# Linking drought strategy to climate of origin in a widespread C4 grass: hydraulic traits, xylem anatomy and stomatal behaviour in *Themeda triandra*

**DOI:** 10.64898/2025.12.03.691378

**Authors:** Vinod Jacob, Chris J. Blackman, Brendan Choat, Brian J. Atwell, Eloise Foo, Alejandro Correa-Lozano, Travis Britton, Emma E. Sumner, Ian J. Wright

## Abstract

- Hydraulic, stomatal and anatomical traits strongly influence drought survival and productivity, but the extent to which these trait-types are coordinated and relate to climate-of-origin in herbaceous species remains poorly understood.
- We quantified hydraulic, stomatal and anatomical traits in eight populations of *Themeda triandra* selected from diverse temperature and precipitation regimes across Australia, grown under common conditions and exposed to drought. Trait – climate-of-origin relationships were quantified using multiple linear mixed models.
- Leaf xylem embolism resistance (P_50_) was unrelated to climate-of-origin precipitation. Instead, plants from drier climates exhibited lower maximum stomatal conductance, earlier stomatal closure during drought and higher stomatal safety margins (the difference between water potential at stomatal closure and P_50_). Plants from warmer climates exhibited greater embolism resistance and higher hydraulic conductance, along with wider vessels and lower vein density, suggesting adaptation to high evaporative demand.
- Our findings highlight two primary hydraulic strategies related to climate-of-origin: 1) conservative water use and drought avoidance in drier climates, and 2) embolism resistance and hydraulic efficiency in warmer climates. These findings provide new insights into the intraspecific differences in traits and adaptive strategies used by C_4_ grasses to persist in contrasting climates.

## INTRODUCTION

Understanding the physiological and anatomical adaptations that underpin plant success in different environments is a primary goal of plant ecology and critical for predicting vegetation responses to climate change (Bjorkman *et al*., 2018). Hydraulic traits, which govern how plants transport and retain water, play a central role in determining plant survival and productivity (Choat *et al*., 2012). A key measure of how plants maintain function under water deficit is the water potential (-MPa) at which 50% of hydraulic conductivity is lost due to embolism, P_50_ (Choat *et al*., 2018). While resistance to xylem embolism defines the lower bounds of water potential that a plant can tolerate before irreversible drought injury occurs, the likelihood of reaching this critical threshold is largely modulated by stomatal regulation (Skelton *et al*., 2015, *Martin-StPaul et al*., 2017). Stomata respond to soil water deficit through gradual closure, minimizing water loss and slowing further declines in xylem water potential. By comparing water potential thresholds of stomatal closure (e.g., 90% closure; g_s90_) and P_50_, a “stomatal safety margin” (SSM) can be defined where SSM = P_50_ – g_s90_. Hydraulic strategies vary widely among plants, ranging from conservative (isohydric; stomata close in advance of xylem failure/high SSM) to risky (anisohydric; stomata remain open even after the onset of embolism/neutral or negative SSM) (Hochberg *et al*., 2018; Hartmann *et al*., 2021). In this study we examined variation in P_50_, g_s90_ and SSM across populations of the widespread C4 grass, *T. triandra*, grown under common conditions. We tested hypothesised relationships between these key traits and climate-of-origin (i.e., the climate from where seeds were sourced), investigating the anatomical basis of hydraulic trait variation, and coordination between hydraulic and photosynthetic traits.

In common-environment studies, hydraulic traits often relate to climate-of-origin (from where seeds or seedlings were sourced), with species from drier environments typically exhibiting more negative P_50_ values, indicating greater resistance to embolism (Bourne *et al*., 2017; Li *et al*., 2018; Volaire *et al*., 2018). Similarly, stomatal closure thresholds (g_s90_), as well as leaf turgor loss point (the water potential at which leaves begin to wilt; Ψ_TLP_), also tends to decline with aridity (Bourne *et al*., 2017; Li *et al*., 2018; Blackman, 2018), such that species from drier sites are able to maintain leaf gas exchange at more negative water potentials during drought. Notably, a global meta-analysis found that stomatal safety margins increase with embolism resistance (Martin-StPaul et al., 2017), indicating that early stomatal closure remains important even in arid-adapted species.

Compared to trait associations with site rainfall, relationships between hydraulic traits and site temperature are less well studied and show inconsistent patterns. While some common garden studies report no relationship between temperature of origin and P_50_ (Li *et al*., 2018; López *et al*., 2021), meta-analyses have reported conflicting trends between temperature and leaf P_50_, ranging from a weak negative relationship (Nardini & Luglio, 2014) to a positive relationship (Huang *et al*., 2024). Such mixed results may not be surprising given that increased embolism resistance is advantageous both in cold environments, where there is increased risk of freeze-thaw embolism, as well as in hot-dry environments, where there is increased risk of embolism during drought (Davis *et al*., 1999; Pittermann & Sperry, 2006; Hartill *et al*., 2023).

Anatomical differences in the water conducting xylem underpin variation in key hydraulic traits, yet their role in modulating variation in hydraulic conductivity and embolism resistance across environmental gradients remains poorly understood. Hydraulic conductivity increases with conduit diameter to the fourth power (Sperry *et al*., 2006) but wider vessels may be more vulnerable to embolism, as proposed by the hydraulic safety-efficiency trade-off (Tyree *et al*., 1994), although empirical evidence for the trade-off is mixed (Gleason et al 2016). Wide vessels are advantageous in warm environments, where high evaporative demand necessitates efficient water transport, but are disadvantageous in dry environments, where the risk of drought-induced embolism increases (Olson *et al*., 2018; Fontes *et al*., 2022). Vessel wall thickness is also closely linked to embolism resistance (Lens *et al*., 2016; Li *et al*., 2016; Volaire *et al*., 2018), possibly via thicker pit membranes (Li et al., 2016; Thonglim et al., 2021), and may also function as an adaptation to freeze-thaw embolism in cooler climates (Davis *et al*., 1999).

Much of our current understanding of plant hydraulic traits and their anatomical basis comes from studies of woody species. By contrast, hydraulic traits in herbaceous plants remain comparatively understudied, and their relationships with climate are rarely explored (but see Ocheltree *et al*., 2016; Volaire *et al*., 2018; Jardine *et al*., 2021). Furthermore, while hydraulic traits have been well described across broad taxonomic scales (e.g., Choat *et al*., 2012; Nardini and Luglio, 2014; Huang *et al*., 2024), within-species variation, especially in herbaceous species, is not well understood. Grasses dominate over 40% of the Earth’s terrestrial surface and are crucial for global agriculture (McSteen & Kellogg, 2022), yet studies of within-species variation in hydraulic and gas exchange traits in grasses from contrasting climates, including the relationship between xylem embolism and stomatal regulation that define different drought response strategies, are lacking.

The traits and processes involved in the survival and recovery of grass species after extreme drought are often complex and multifaceted (Craine *et al*., 2013; Hoover *et al*., 2014). Like woody species, perennial grasses from arid environments may exhibit drought tolerance (embolism resistance) or drought avoidance (early loss of turgor and stomatal closure) strategies, but they can also employ alternative mechanisms such as aboveground senescence and/or dormancy to survive drought (Volaire *et al*., 2009; Zwicke *et al*., 2015; Balachowski *et al*., 2016; Keep *et al*., 2021). *Themeda triandra Forssk*. is a perennial C_4_ tussock grass in the Andropogoneae tribe (which includes major crops like sorghum, maize, and sugarcane) and is the most widespread plant species in Australia (Gallagher, 2016), occurring across all biomes from semi-arid savannas and seasonally wet tropics to sub-alpine and cool temperate regions (Dunning *et al*., 2017). The exceptional climatic breadth and ecological dominance of this species make it ideal for examining how grasses have evolved diverse strategies to cope with drought and to investigate how variation in hydraulic and other physiological traits are driven by climate adaptation.

In this study *T. triandra* plants were grown from seed sourced from eight populations from sites across the Australian continent strongly contrasting in rainfall and temperature. We measured a range of key hydraulic, photosynthetic and xylem anatomical traits in well-watered plants before characterising their stomatal response to drought. We tested hypothesised relationships between key traits and climate-of-origin rainfall and temperature (see below). We also explored various trait– trait relationships, including those related to the proposed hydraulic safety–efficiency trade-off (Hacke *et al*., 2006; Gleason *et al*., 2016; Ocheltree *et al*., 2016) and the coordination between leaf hydraulics and leaf gas exchange . By investigating hydraulic, anatomical and gas exchange traits across multiple *T. triandra* populations grown under common conditions, this study aims to identify distinct trait-climate associations based on genetic variation without the influence of environmentally-induced plasticity (Jankowski *et al*., 2019; Huxman *et al*., 2022; Schwinning *et al*., 2022). Understanding these within-species patterns in a wild Andropogoneae grass may provide insights for predicting shifts in native grassland composition under climate change and avenues for enhancing drought resilience in related crop species.

Our work tested a number of hypotheses based on fundamental principles of plant hydraulics, and previous empirical trends reported in the literature:

### Hypotheses

1. Plants from drier environments exhibit more negative values for P_50_, Ψ_TLP_, and g_s90_ and have smaller, thicker-walled vessels and higher vein density.
2. Plants from warmer sites exhibit higher hydraulic conductance (K_leaf_), linked to wider vessels, enabling more efficient water transport and sustained leaf function under high evaporative demand.
3. Plants with more negative P_50_ have smaller vessels and thicker xylem vessel walls accompanied by lower K_leaf_, in accordance with an assumed hydraulic safety-efficiency trade-off.

In addition, we expected traits related to rapid resource capture (A_max_, K_leaf_, and g_smax_) to covary positively among themselves but negatively with traits related to stress tolerance (P_50_, Ψ_TLP_, and g_s90_), reflecting a trade-off between distinct adaptive strategies.

## Methods

### Plant Growth

We grew 10 – 12 replicates of eight populations of *T. triandra*, sourced from diverse climatic regions across Australia (Figure 1, Table S2; summer temperatures ranging from 16.3 – 32.1°C and summer precipitation ranging from 75 – 435 mm), in a climate-controlled glasshouse at the Hawkesbury Institute for the Environment, Western Sydney University, Richmond, NSW. Plants were grown from seeds germinated in seed trays. After emergence they were transplanted into 8-litre pots filled with sandy-loam soil (pH ≈ 5.6) collected from the PACE field facility at Western Sydney University (Churchill *et al*., 2022). The soil in each pot was supplemented with 10 g of slow-release native-plant fertiliser (Osmocote® Native). Seedlings subsequently received a half-strength application of Yates Thrive® All-Purpose fertiliser (N:P: K 25:5:8), applied weekly for the first four weeks, to prevent nutrient limitation during establishment.

**FIGURE 1.**
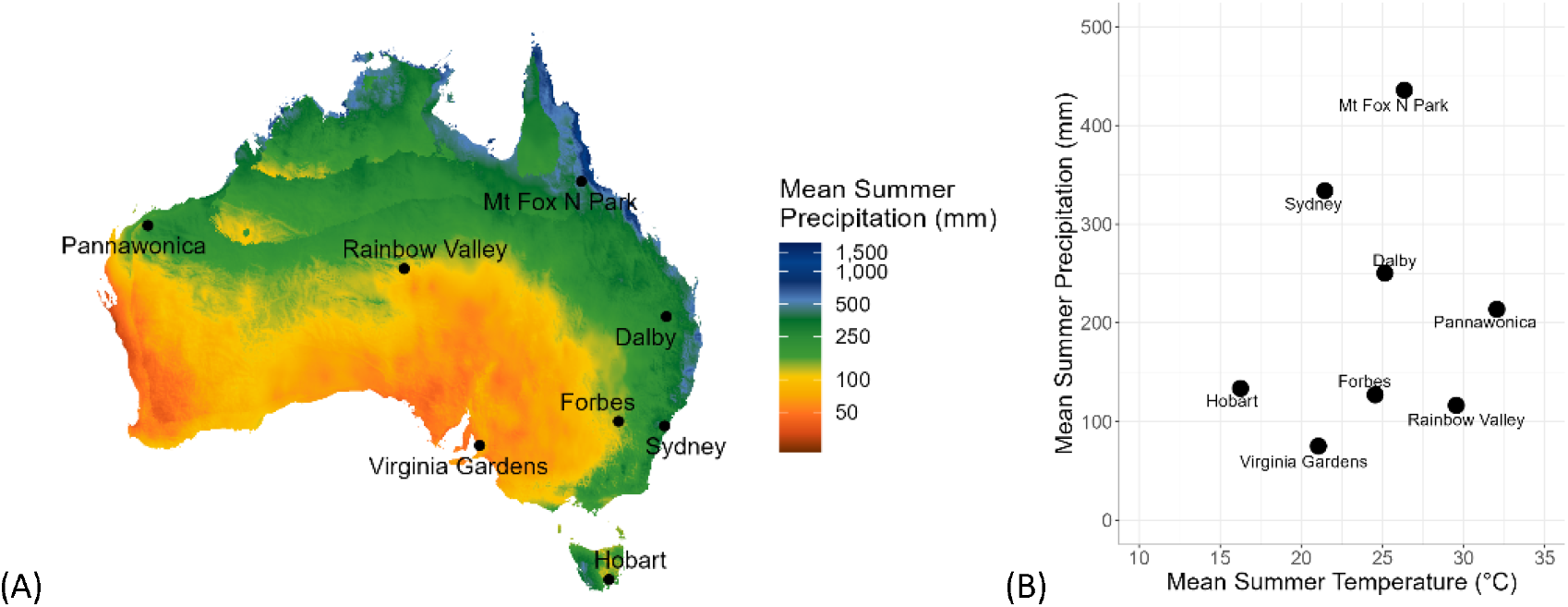
Climate-of-origin of populations used in this study. (A) Mean summer precipitation; mm) across Australia; populations represented by black dots (•). (B) Scatter plot illustrating the two key climatic predictors, with summer temperature (°C) on the x-axis and mean summer precipitation on the y-axis.

All pots were maintained at 100% field capacity, using an automated irrigation system, for the first two months of growth. The glasshouse was maintained at 30/20°C (day/night) temperatures with ambient CO_2_ levels (∼420 ppm) and a 14-h light period. Natural light was supplemented throughout the light period with full spectrum LED lighting, adding 700 μmol photons m^-2^ s^-1^, with maximum midday levels reaching ∼1500 μmol photons m^-2^ s^-1^. Following this initial growth period, all plants underwent a “drought-hardening” pre-treatment, in which watering was withheld for 4 – 5 days, inducing visible leaf wilting. After this period of water deficit, plants were rewatered to 100% field capacity and maintained under these conditions for 4 – 6 days, ensuring full rehydration (predawn leaf water potential ≈ 0). All plants were maintained at 100% field capacity for an additional two weeks before being allocated to two experimental groups: 1) “Steady-State Trait Characterisation” plants were used to measure key hydraulic, physiological and anatomical traits, and 2) “Dynamic Dry-Down Assay” plants were subjected to a progressive dry-down to assess stomatal responses to desiccation.

### Steady-State Trait Characterisation

#### Leaf xylem vulnerability measurements

The optical vulnerability (OV) technique (Brodribb *et al*., 2016) was used to visualise embolism events and estimate vulnerability thresholds in the leaves of 4 – 5 replicates for each population. Whole individual plants (including root systems) were carefully removed from pots early in the morning and roots were washed to remove any soil. Excised plants were placed in a bucket containing a shallow layer of water and covered with black plastic bags to ensure that they remained well hydrated, so embolism due to dehydration did not occur before OV measurements began. Plants were transported immediately to the laboratory for OV measurements, where a fully expanded leaf from each individual was positioned between two plastic polarising films and placed into a CaviCam clamp. The CaviCam consists of a Raspberry Pi single board computer (Raspberry Pi Foundation, http://www.raspberrypi.org) which controls and stores images produced by an 8-Mp camera magnified by a ×20 lens. Leaf samples were illuminated from below using six light-emitting diodes (LEDs) delivering ∼200 μmol quanta m^−2^ s^−1^ to the leaf and providing refracted light from the sample surface to the camera. All components were housed within a 3D-printed clamp (further details of the clamps are provided at https://github.com/OpenSourceOV).

Leaf water potential was concurrently measured on the nearest neighbouring leaf using a stem psychrometer (ICT PSY Armidale, NSW, Australia) (Johnson *et al*., 2018). The psychrometer was automated to take water potential readings every 10 minutes, and the accuracy of the psychrometer was verified for a subset of individuals using a Scholander pressure chamber (PMS Instruments) (Figure S1); leaves of comparable size were used for all individuals for consistency. Both water potential and xylem images were recorded until each plant had dehydrated completely (i.e. embolism events had ceased for >12 hours).

Images of xylem embolism were processed using ImageJ using the instructions at https://github.com/OpenSourceOV/image-processing-instructions/blob/master/instructions.md. Differences between neighbouring images in each sequence were used to highlight embolism events. Rapid changes in pixel intensity associated with embolism are distinguishable from other changes in the leaf during dehydration (Johnson *et al*. 2018). Threshold analyses were used to remove noise related to leaf movement, in order to produce a set of images that contain only the pixels associated with embolism events. These data were used to determine the percentage of embolism using the following equation.

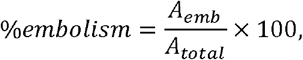

Where A_emb_ is the cumulative area of embolised xylem at a given time and A_total_ is the total area of embolised xylem at the end of sample dehydration. Finally, the water potential data were combined with the total cumulative pixel count to create a vulnerability curve for each species.

#### Maximum rates of leaf gas exchange and leaf hydraulic conductance

Steady state measurements of net photosynthesis (A_sat_ ; μmol m^−2^ s^−1^), stomatal conductance (g_s_ ; mol m^−2^ s^−1^) and transpiration (*E*; mmol m^−2^ s^−1^) were conducted on 4 – 5 replicates of each population between 09:00 and 12:00 on recently fully expanded leaves using a portable open, infra-red gas exchange system (LI-6800, Li-Cor, Lincoln, NE, USA). Measurements were conducted at a saturating photosynthetic photon-flux density (PPFD) of 1800 mmol m^-2^ s^-1^ and a chamber reference [CO_2_] (C_a_) of 420 μmol CO_2_ mol^−1^ at average daily mid-day temperatures (30°C). Leaf-to-air vapour pressure deficit levels were maintained at roughly 1.5 – 2.0 kPa. Each leaf was allowed ∼5 mins to equilibrate before measurements were taken (Ghannoum *et al*. 2010). Well-watered control pots from experiment two were also measured daily until the end of the experiment and *A*_*sat*_ and g_s_ values were averaged across multiple timepoint to calculate g_smax_ and A_max_.

Maximum leaf hydraulic conductance (K_leaf_) was measured using the “relaxation by rehydration” method (Brodribb & Holbrook, 2003). A pair of adjacent leaves were selected based on comparable size and condition from well-watered individuals (either two leaves from the same tiller or adjacent leaves). The initial leaf water potential (Ψ_0_) was measured on the first leaf, using a pressure chamber, and the second leaf was partially submerged in filtered, de-ionized water, and recut at the base and rehydrated for 30 seconds. The apical section of the leaf remained out of the water, ensuring that rehydration occurred only through the cut xylem. After rehydration, the second leaf was re-cut above the water line and placed in a pressure chamber to obtain the final rehydrated leaf water potential (Ψ_f_). Leaf hydraulic conductivity (K_leaf_) was calculated using the following equation.

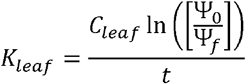

where C_leaf_ is the bulk leaf capacitance before turgor loss, normalized by leaf area, t is the duration of rehydration, and Ψ_0_ and Ψ_f_ are the initial and final leaf water potentials, respectively.

#### Pressure Volume Curves

Pressure-volume curves were conducted on multiple leaves from 4 – 5 individuals from each population to determine turgor loss point (ΨTLP) and C_leaf_. Leaves were sampled at dusk and placed in opaque plastic bags, with the distal end of each leaf placed under water and left to rehydrate overnight. Rehydrated leaves were photographed, and images were analysed using ImageJ to calculate leaf area. Leaf fresh weight (FW) and Ψ_leaf_ were then periodically measured during slow desiccation, until Ψ_leaf_ reached -3 MPa. Sample leaves were then oven dried at 70°C for a minimum of 48 hr to obtain dry weight (DW). Ψ_TLP_ was calculated by plotting leaf relative water content (RWCleaf, %) against the inverse of Ψ_leaf_, and Ψ_TLP_ was determined as the point where the line became non-linear. Area-based leaf-specific capacity at full turgor (C_leaf_) and leaf osmotic potential at full turgor (Ψ_osm_) were calculated for each individual leaf following standard pressure-volume curve analysis methods (Turner, 1988; Sack et al., 2011) and averaged across replicate leaves of the same plant.

#### Analyses of xylem characters using light microscopy

Leaf samples from the same individuals used for leaf hydraulic vulnerability measurements were collected and immediately fixed in a 5% formaldehyde, 5% acetic acid, and 70% ethanol solution (FAA) solution for 48 hours to preserve tissue structure. Following fixation, samples were gradually dehydrated through an ethanol series up to 70% for long-term storage before embedding. Fixed samples were embedded in Optimal Cutting Temperature compound (Tissue-Tek® O.C.T. Compound, Sakura Finetek, USA) and sectioned at -25°C and 6 µm thickness using a cryostat (HM525 NX, Epredia). Sections were transferred by gentle pressure contact onto 1.3% Fol’s chrom-alum gelatine subbed glass slides (Kiernan, 1999). For staining, sections on slides were first submerged in 1% Safranin O for 10 minutes, to stain lignified tissues. Samples were then dehydrated in 70% ethanol and counterstaining was performed with 0.5% Fast Green (in absolute ethanol). Finally, sections were cleared in xylene and dry specimens were mounted permanently with Eukitt mounting medium (Sigma-Aldrich).

Images from each population were captured using an Axiocam 503 colour module (Zeiss) attached to an Axioscope 5 (Zeiss) microscope under a phase contrast setup. Leaf anatomical traits were quantified via image analysis using Fiji, ImageJ2 (ImageJ v. 1.54f). Measured traits included vein density and diameter for both major and minor veins, and metaxylem vessel wall thickness for major veins. For each replicate, two measurements were taken and the mean reported. Vein diameter was quantified as the tangential diameter of the two largest vessels, measured at the widest point of the lumen (excluding cell walls), and expressed as arithmetic means. Vessel wall thickness was measured on metaxylem vessels in major veins. Vein density was determined by counting individual major and minor veins per leaf section and normalizing by leaf section length (veins cm^−1^), determined using a white reference line along the blade margin. Further methodological details, measurement protocols and representative images, are provided in Table S4.

#### Dynamic Dry-Down Assay

One replicate pot from each population was maintained at 100% field capacity to serve as the well-watered control while four other replicates of each population were subjected to a prolonged, severe dry-down by completely withholding irrigation. Soil moisture was measured daily using a handheld soil water reflectometer (HydroSense II CS659, Campbell Scientific). Predawn (Ψ_PD_) and midday (Ψ_MD_) leaf water potentials were also measured daily, using a Scholander pressure bomb (PMS Instruments, Albany, OR, USA) fitted with a grass compression gland and base, on two to three leaves per individual to track plant water status. Concurrently, daily measurements of light-saturated photosynthesis (A_*sat*_; μmol m^−2^ s^−1^), stomatal conductance (g_s_ ; mol m^−2^ s^−1^) and transpiration (*E*; mmol m^−2^ s^−1^) were taken on the same leaves. These data were used to derive the leaf water potential at 90% stomatal closure (g_s90_). Steady state measurements of *A*_sat_, g_s_ and *E* were conducted between 09:00 and 12:00 on recently fully expanded leaves using the LI-6800 (described above). Measurements were conducted at a saturating PPFD of 1800 mmol m^-2^ s^-1^ with a chamber reference [CO_2_] (C_a_) of 420 μmol CO_2_ mol^−1^ and at average daily midday temperatures (30°C). Leaf-to-air vapour pressure deficit levels were maintained at roughly 1.5–2.0 kPa. Each leaf was allowed ∼5 mins to equilibrate before measurements were taken. Measurements were conducted daily on the dehydrating pots (on 2 – 3 leaves from each individual plant per day) until stomatal closure occurred (i.e. g_s_ was reduced by > 90%). All parameters were also measured on well-watered control pots until the end of the experiment.

#### Climate data

*T. triandra* occurs across a variety of climate zones in Australia. To identify the most appropriate climate-of-origin descriptors for this comparative study, ten climatic variables capturing temperature and precipitation means, seasonality, and extremes for each of the eight source locations were extracted from CHELSA gridded datasets (Karger *et al*., 2017), which report long-term (1981 – 2010) interpolated climate averages at a spatial resolution of ∼ 1km at the equator. Principal components analysis was used to identify the major tendencies across these ten variables, with varimax rotation applied to improve interpretability (aligning climate vectors more clearly with individual factors rather than composite mixtures). The first rotated axis reflected a temperature–seasonality gradient (driven by temperature range and mean summer temperature). The second axis captured a precipitation gradient (driven by summer and annual precipitation). From these analyses we selected *summer temperature* and *summer precipitation* as the key climate variables to be used in analysis of trait – climate relations. These two variables had the strongest loadings (Table S3) on their respective axes and have clear biological relevance given this perennial species exhibits peak growth during the warmer summer months, especially in colder regions where its growth is restricted to this period (Snyman *et al*., 2013; Churchill *et al*., 2022). Summer temperature and summer precipitation vary almost independently across our eight populations (R^2^ = 0.03, P = 0.698), allowing their use in subsequent models. Note, we used statistical approaches (described below) that simultaneously considered the effects of two climate variables, allowing us to quantify trait relationships with summer temperature *with variation in summer rainfall held constant*, and vice versa.

#### Statistical Analysis

Leaf hydraulic vulnerability curves were fitted using the “fitplc” function from the “fitplc” package (Duursma & Choat, 2017), using Weibull models where they converged and sigmoidal models when curves exhibited abrupt, steep declines in conductance that caused non-linear optimisation to break down. Individual vulnerability curves were generated for each replicate plant and the leaf water potentials causing 12% (P_12_) and 50% (P_50_) xylem embolism were quantified. Stomatal conductance values were plotted against leaf water potential and fitted with a weighted polynomial regression, using the “*fitcond*” function within the “*fitplc*” package, to extract leaf water potential at stomatal closure (g_s90_). Stomatal safety margins (MPa) were calculated as the difference between the Ψ_MD_ at which stomata were 90% closed (g_s90_) and P_50_ (SSM).

To assess the relationship between plant functional traits and the two climate variables, we used linear mixed-effects models fitted with the “lme4” package in R. The model structure included a response variable (trait of interest) as a function of climate predictors. To incorporate the repeated sampling structure of the data, the site of origin for each population was specified as a random effect:

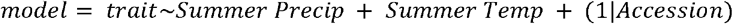

A corresponding fixed-effects linear model was also fit without the random effect for visualization of partial residuals. We also tested for an interaction between summer precipitation and summer temperature for each trait but found no significant interactions, so we retained an additive model structure. The effect of each climate predictor on the response trait was visualized using the “visreg” package, extracting model residuals to generate partial residual plots. This approach allows for the visualization of the independent effects of precipitation and temperature on the trait of interest while accounting for the influence of other variables in the model. P-values for fixed effects were quantified using the “lmerTest” package.

To assess the robustness of our findings, we also conducted bivariate analyses of each climate variable independently. This approach allowed us to examine whether incorporating both precipitation and temperature in the mixed model influenced our results by accounting for covariation between these climate factors. Additionally, bivariate analysis strengthens our results by providing a complementary perspective on trait-climate relationships, helping us determine which correlations remain consistent across different analytical approaches. Moreover, we included an annual moisture index (calculated as precipitation/potential evapotranspiration) in our bivariate analysis, this being a widely used, an integrated measure of site water availability.

## Results

Leaf hydraulic conductance (K_leaf_) varied markedly among *T. triandra* populations, ranging from an average of 2.53 mmol m^−2^ s^−1^ MPa^−1^ at Sydney to 11.10 mmol m^−2^ s^−1^ MPa^−1^ at Virginia (Table. S5). Xylem embolism resistance (P_50_) also varied clearly among populations, ranging from -2.41 MPa in plants from Hobart (southern Australia) to -5.13 MPa (Fig. S2) in plants from Pannawonica (north-western Australia). Traits governing plant water status showed less variation across populations, with mean turgor loss point (Ψ_TLP_) ranging from -1.17 to -1.86 MPa and stomatal closure (g_s90_) ranging from -3.93 to -5.44 MPa. Stomatal safety margins (SSM = P_50_ - g_s90_) ranged from +0.75 MPa; Forbes) to -1.54 MPa; Mt Fox National Park)., with positive and negative values indicating that stomatal closure occurs before and after the onset of 50% embolism, respectively.

### Trait relationships with climate-of-origin

We predicted (Hypothesis 1) that populations from drier sites would be more resistant to water stress, displaying more negative P_50_ and Ψ_TLP_ values than those from wetter sites. Instead, we found that P_50_ and precipitation were unrelated (Fig. 2A), while drier-site populations exhibited less negative Ψ_TLP_ (Fig. 2C). These findings were supported by both the LMM and bivariate analyses (Table S1). Drier-site populations exhibited a suite of drought-avoidance traits, including stomatal closure at less negative water potentials (g_s90_) and wider stomatal safety margins (SSM) (Fig. 2G). K_leaf_ was significantly higher in drier-site populations (Fig. 2I). Neither of the SSM or K_leaf_ trends were observed in bivariate analyses (Table S1).

**Figure 2.**
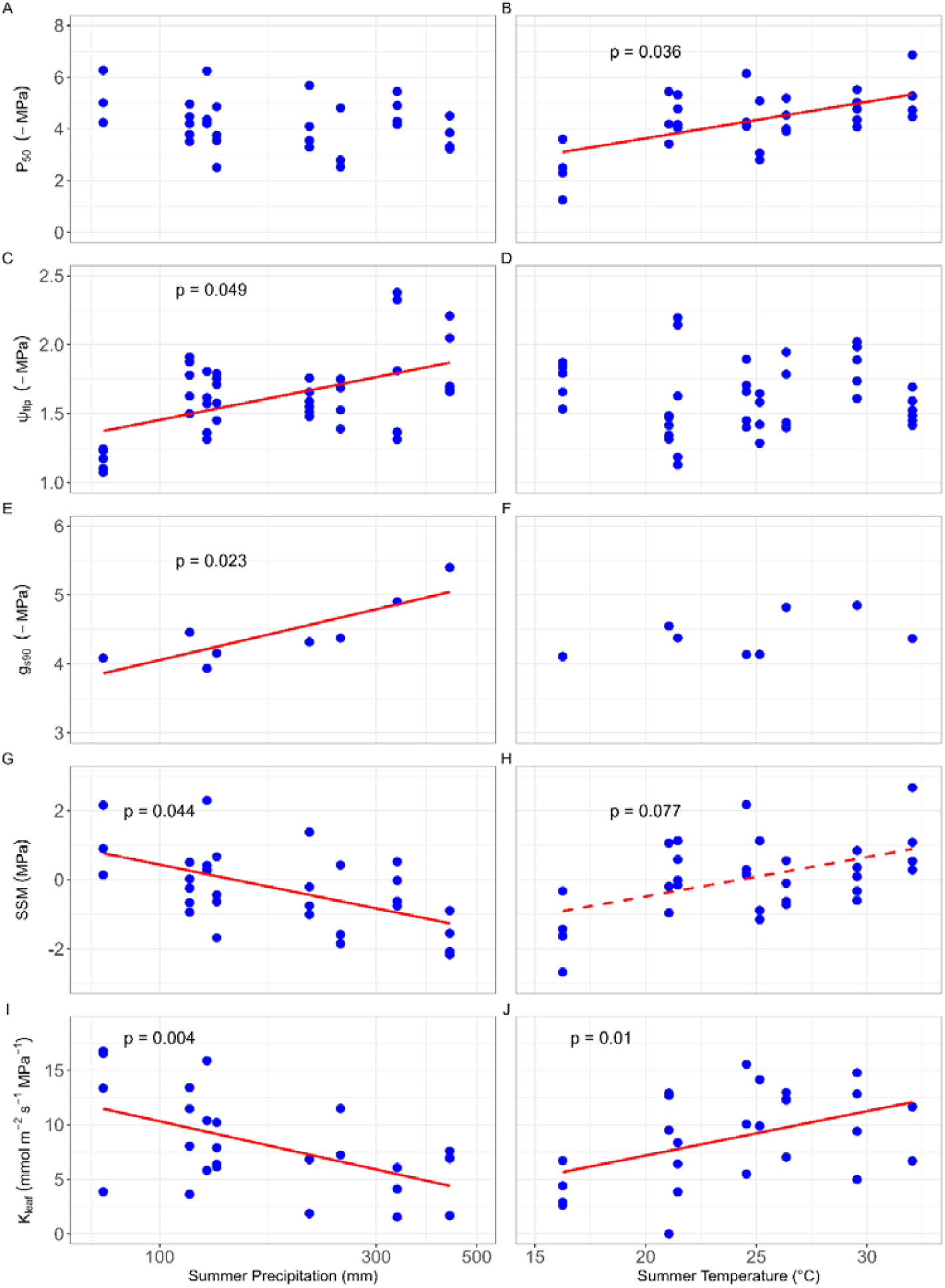
Partial residual plots showing the independent relationships between climate-of-origin and hydraulic traits across populations of *T. triandra*. Relationships between mean summer precipitation (A, C, E, G, I) and mean summer temperature (B, D, F, H, J) with (A, B) water potential at 50% xylem embolism (P_50_), (C, D) water potential at leaf turgor loss (Ψ_TLP_), (E, F) water potential at 90% stomatal closure (g_s90_), (G, H) stomatal safety margin (SSM, Δ g_s90_ & P_50_), and (I, J) leaf hydraulic conductance (K_leaf_). Fitted lines are derived from linear mixed-effects models using mean data from individual plants, with population as a random effect. Partial residuals were obtained from a corresponding multiple regression model, without the random effect, to visualize the independent effects of each climate variable. Solid red lines indicate significant linear regression fits (*p* < 0.05), while dashed red lines denote marginally significant relationships (*p* < 0.1). Mean summer precipitation was log-transformed to standardize coefficients and back-transformed for visualization.

Using LMM analysis, populations originating from warmer climates exhibited greater drought tolerance, as indexed by more negative P_50_ (Fig. 2B). However, Ψ_TLP_ (Fig. 2D) and g_s90_ (Fig. 2F) were unrelated to climate-of-origin temperature. These tendencies were also observed in bivariate trait– temperature analyses (Table S1). The positive relationship between temperature and P_50_ indicated a tendency for plants from warmer climates to exhibit higher SSMs than plants from cooler climates (Fig. 2H), although this was observed in the LMM analysis only. In support of Hypothesis 2, warm-origin plants had significantly higher K_leaf_ than cool-climate plants (Fig. 2J), if in the LMM analysis only (Table S1).

Moisture Index (the annual balance between precipitation and potential evapotranspiration) at the climate of origin had little relationship to hydraulic traits except that plants from more arid environments exhibited marginally higher hydraulic conductance than those from mesic sites (P = 0.067; Table S1).

Counter to expectations (H3), neither vessel diameter (in major or minor veins) nor vessel wall thickness varied with climate-of-origin rainfall, whether considered simultaneously with temperature variation (LMMs: Figs 3A, E, I) or in bivariate analyses (Table S1). Vessel density (number per cross-sectional area) was also unrelated to precipitation (Fig. 3C, G; Table S1). Relationships with site temperature differed between vein types: major vein traits were unrelated to temperature (Fig. 3B, D; Table S1), whereas minor vein traits showed distinct temperature-driven trends. Plants from warmer sites had wider minor vein vessels (Fig. 3F) and reduced minor vein density compared to cooler sites (Fig. 3H). The same trends were also evident in bivariate trait– climate analyses (Table S1). Moisture Index was not correlated with vessel diameter (both minor and major veins) but plants from more arid sites had lower minor vein density than those from more mesic sites (P = 0.049 Table S1).

**Figure 3:**
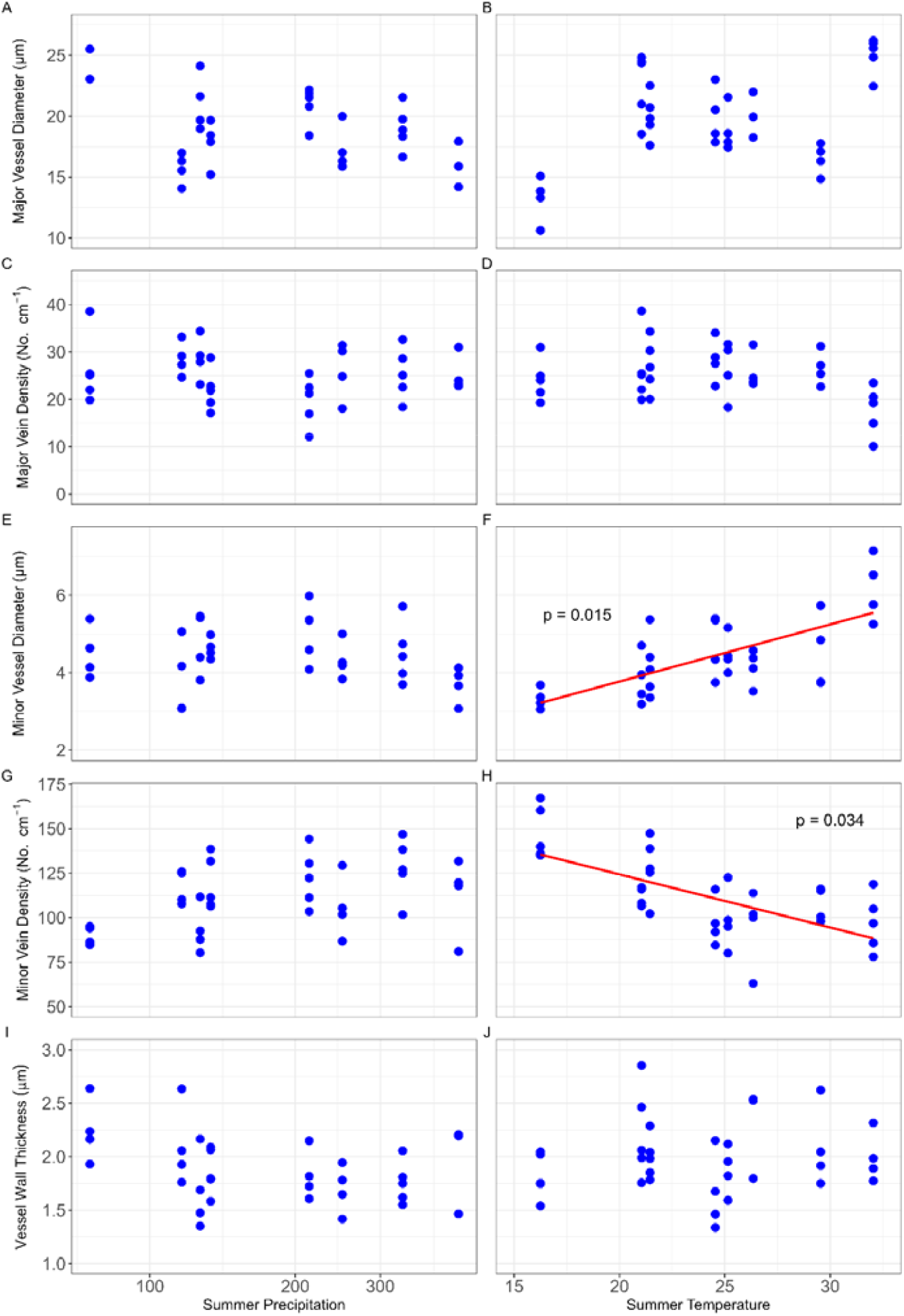
Partial residual plots showing the independent relationships between climate-of-origin and xylem anatomical traits across populations of *T. triandra*. Relationships between mean summer precipitation (A, C, E, G) and mean summer temperature (B, D, F, H) with (A, B) major vessel diameter, (C, D) major vein density, (E, F) minor vessel diameter and (G, H) minor vein density. Fitted lines are derived from linear mixed-effects models using mean data from individual plants, with population as a random effect. Partial residuals were obtained from a corresponding multiple regression model, without the random effect, to visualize the independent effects of each climate variable. Solid red lines indicate significant linear regression fits (*p* < 0.05). Mean summer precipitation was log-transformed to standardize coefficients and back-transformed for visualization.

In LMMs, g_smax_ was significantly higher in plants from wetter environments (Fig. 4C) and marginally higher in plants from colder sites (Fig 4D), however, A_max_ was not significantly related to climate-of-origin rainfall or temperature (Fig. 4A, B). These relationships were consistent in bivariate trait-climate analyses (Table S1).

**Figure 4.**
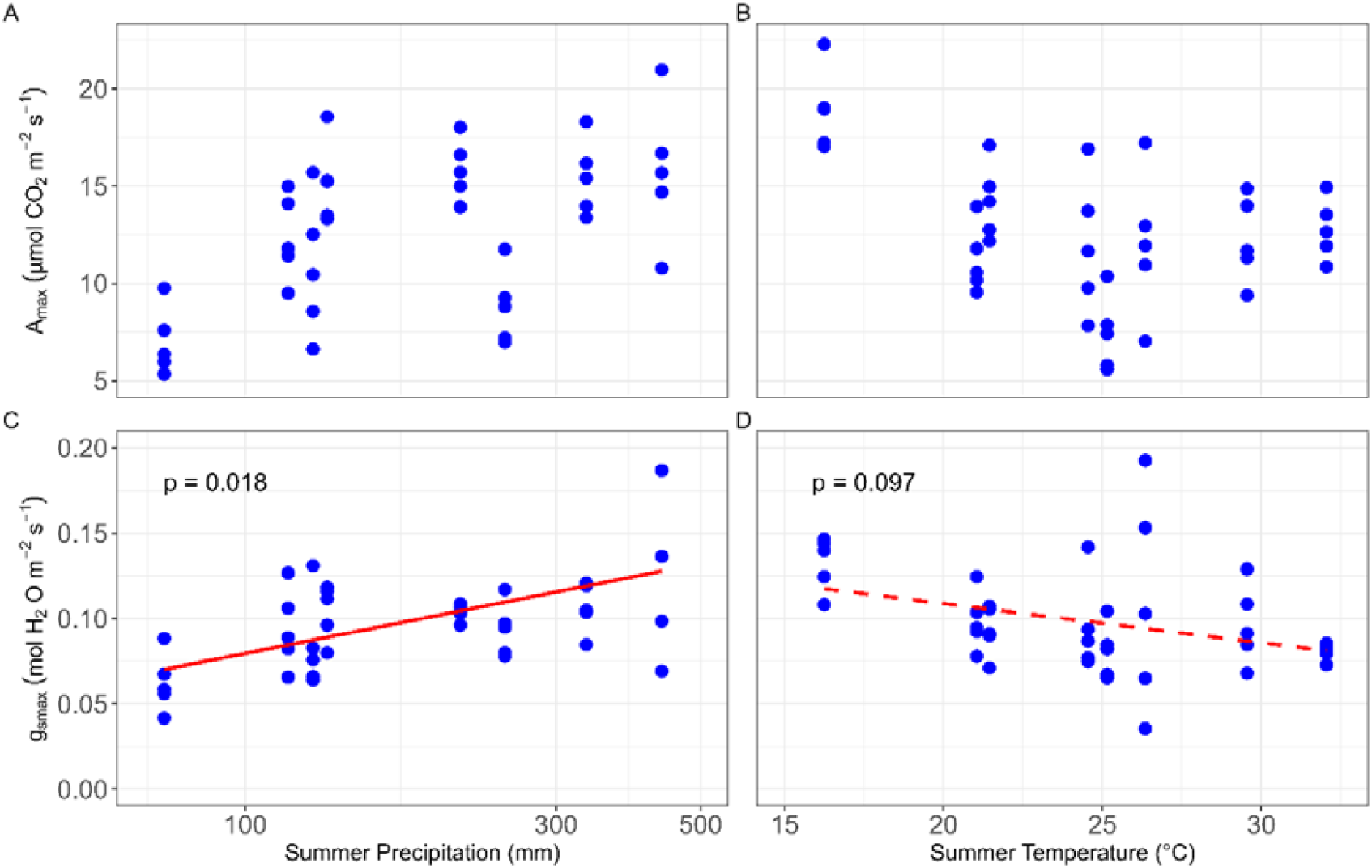
Partial residual plots showing the independent relationships between climate-of-origin and leaf gas exchange traits across populations of *T. triandra*. Relationships between mean summer precipitation (A, C) and mean summer temperature (B, D) with (A, B) maximum rates of light saturated photosynthesis (A_max_), (C, D) and maximum rates of stomatal conductance (g_smax_). Fitted lines are derived from linear mixed-effects models using mean data from individual plants, with population as a random effect. Partial residuals were obtained from a corresponding multiple regression model, without the random effect, to visualize the independent effects of each climate variable. Solid red lines indicate significant linear regression fits (*p* < 0.05), while dashed red lines denote marginally significant relationships (*p* < 0.1). Mean summer precipitation was log-transformed to standardize coefficients and back-transformed for visualization.

### Coordination among hydraulic, anatomical, and photosynthetic traits

The expected trade-off (Hypothesis 3) between hydraulic safety (P_50_) and hydraulic efficiency (K_leaf_) was not supported (Fig. 5A). Variation in P_50_ was also unrelated to A_max_ and g_smax_ (Fig. 5C, D). Notably, populations with lower Ψ_TLP_ typically had lower g_smax_ (R^2^ = 0.55, P < 0.036; Fig. 5H) and marginally lower A_max_ (R^2^ = 0.45, P = 0.067; Fig. 5I). Contrary to expectations (Hypothesis 3), plants that were more resistant to embolism (more negative P_50_) also had wider major vein xylem vessels (R^2^ = 0.61, P = 0.023; Fig. 6A) and wider minor vein xylem vessels (R^2^ = 0.52, P = 0.043; Fig. 6C). Plants with more negative P_50_ also exhibited lower minor vein density (R^2^ = 0.48, *P* = 0.056; Fig. 6D), but there was no relationship with major vein density (Fig. 6B). Vessel wall thickness was unrelated to P_50_ (Fig. 6E). Variation in hydraulic conductivity (K_leaf_) was largely unrelated to xylem anatomy, except that plants with lower minor vein density had higher K_leaf_ (R^2^ = 0.79, P = 0.003; Fig. 6I). We found no evidence of coordination between xylem embolism resistance (P_50_) and stomatal closure (g_s90_; Fig. 7A, C, E) or turgor loss point (Ψ_TLP_; Fig. 7B, D). However, Ψ_TLP_ exhibited a significant positive relationship with g_s90_ (Fig. 7E), consistent with the role of turgor loss in constraining stomatal closure during dehydration. K_leaf_ was strongly negatively related to both A_max_ (Fig. 8A) and g_smax_ (Fig. 8B), contrasting with a simple supply–demand coupling between hydraulic conductance and gas exchange capacity in this species.

**Figure 5:**
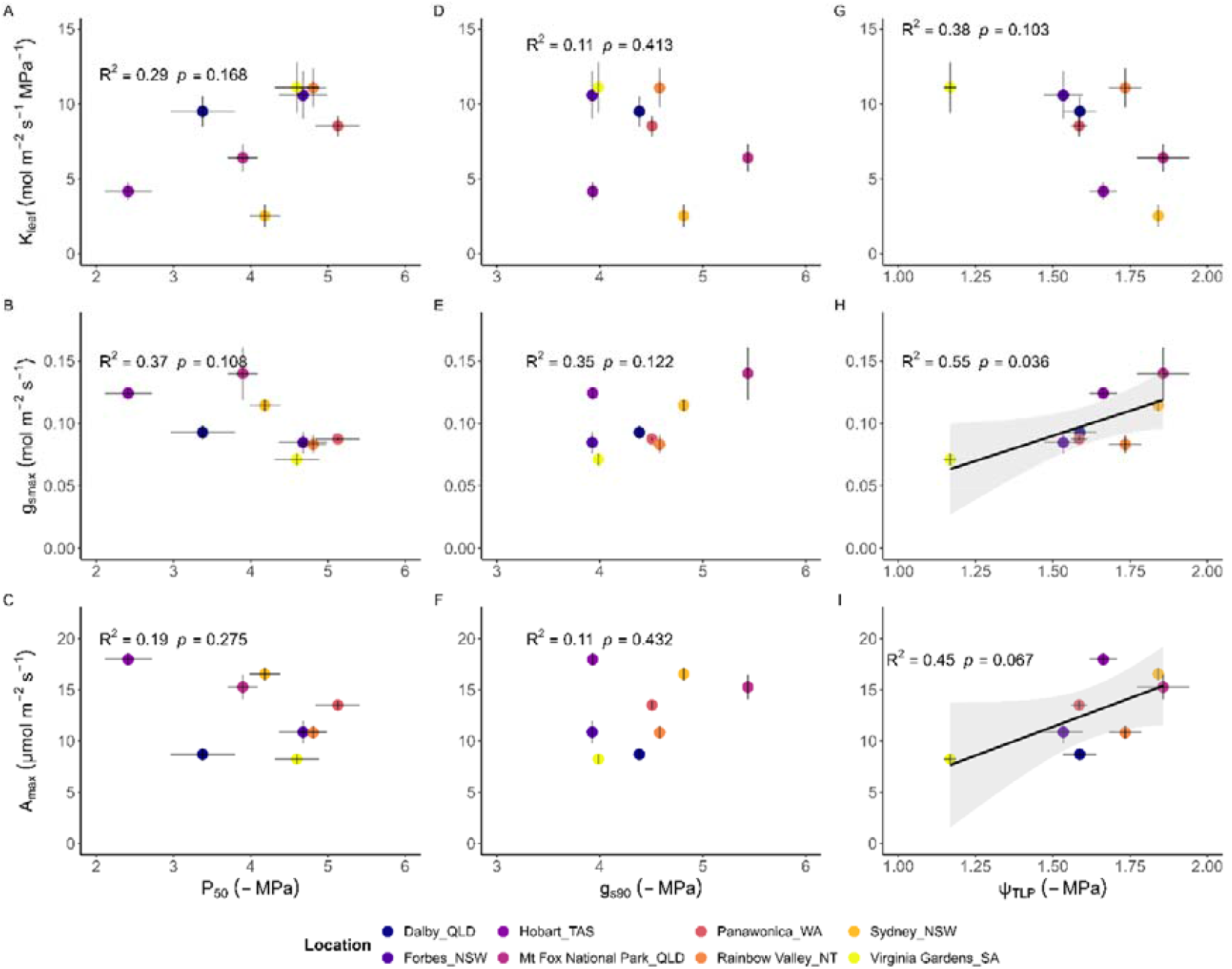
Linear regressions examining relationships between *safety* and *efficiency* traits across populations of *T. triandra*. Correlation between xylem embolism resistance (P_50_) and (A) hydraulic conductivity (K_leaf_), (B) maximum stomatal conductance (g_smax_) and (C) maximum photosynthetic rate (A_max_). Relationships between xylem water potential at 90% stomatal closure (g_s90_) and (D) K_leaf_, (E) g_smax_ and (F) A_max_. Relationship between leaf water potential at turgor loss (Ψ_TLP_) and (G) K_leaf_, (H) g_smax_ and (I) A_max_. Solid lines and shaded 95% confidence bands are displayed only for statistically significant relationships (P < 0.1). Adjusted R^2^ values and P-values (derived from Pearson correlations, equivalent to simple OLS tests) are likewise displayed only when p < 0.1. Error bars denote ± standard error of the mean; colours denote different populations.

**Figure 6:**
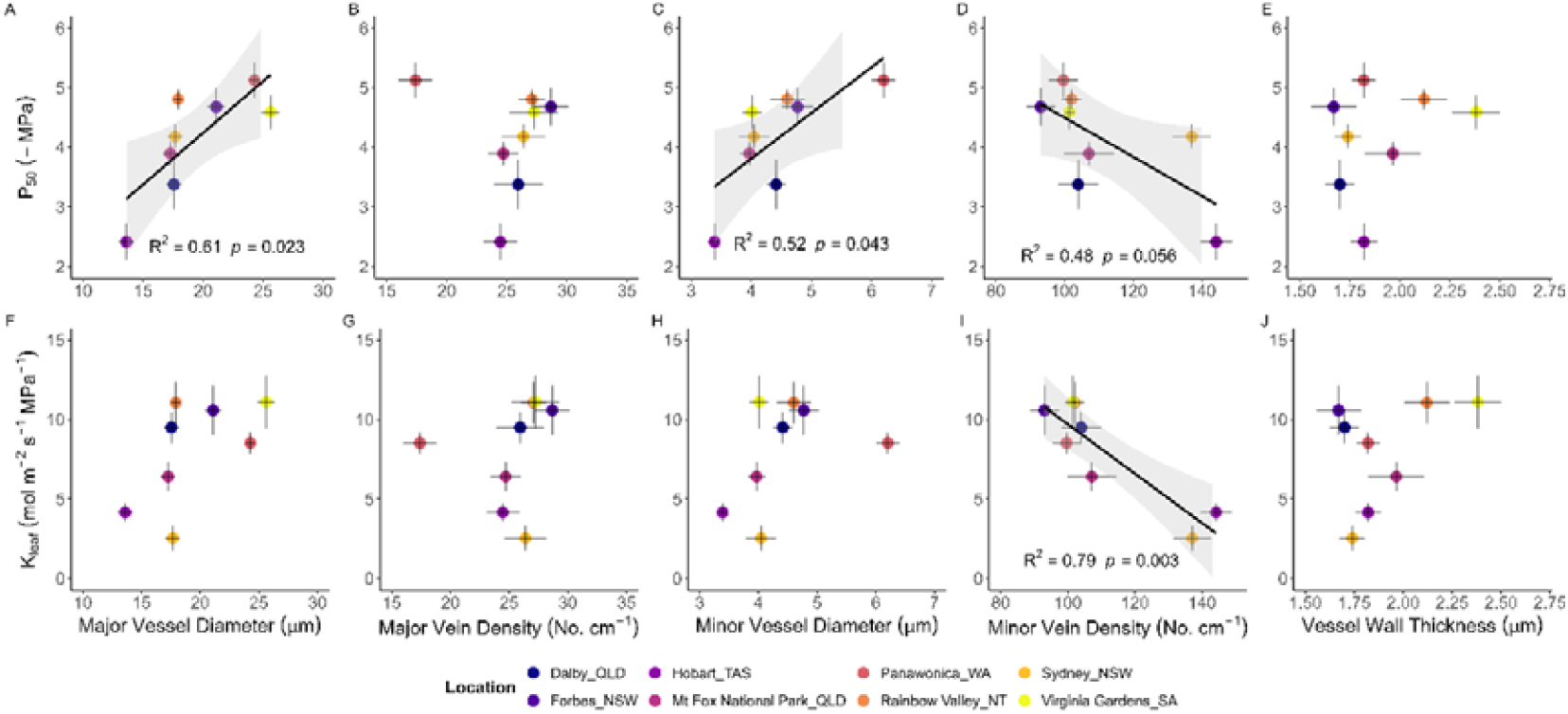
Linear regressions examining relationships between hydraulic traits and xylem anatomy across populations of *T. triandra*. (A–E) Correlations between xylem embolism resistance (P_50_), (F– J) leaf hydraulic conductance (K_leaf_) and major vessel diameter (A, F), major vessel density, (B, G) minor vessel diameter (C, H), minor vein density (D, I), and major vessel wall thickness (E, J). Solid lines and shaded 95% confidence bands are displayed only for statistically significant relationships (P < 0.1). Adjusted R^2^ values and P-values (derived from Pearson correlations, equivalent to simple OLS tests) are likewise displayed only when p < 0.1. Error bars denote ± standard error of the mean; colours denote different populations.

**Figure 7.**
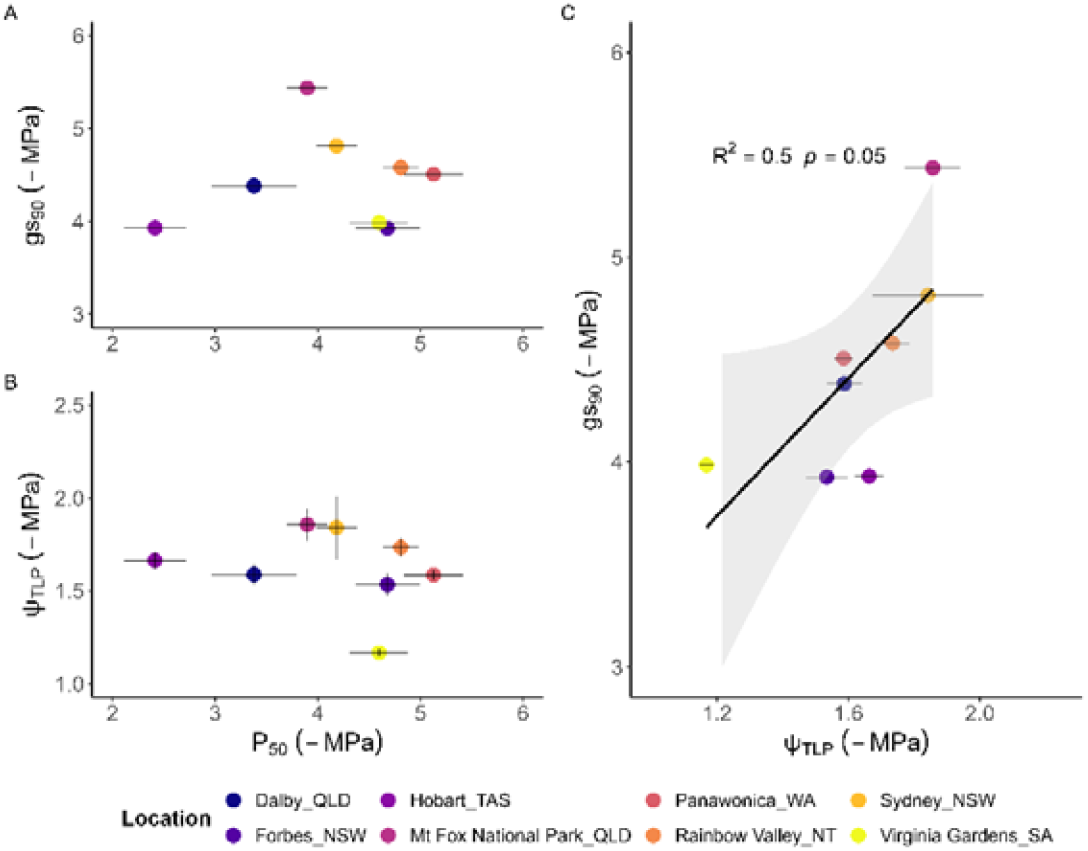
Linear regressions examining relationships between hydraulic vulnerability and water potential at turgor loss and stomatal closure across *T. triandra* populations. Correlations between water potential at 50% xylem embolism (P_50_) and (A) water potential at 90% stomatal closure (g_s90_) and (B) leaf water potential at turgor loss (Ψ_TLP_). Correlation between Ψ_TLP_ and (E) g_s90_ . Solid lines and shaded 95% confidence bands are displayed only for statistically significant relationships (P < 0.1). Adjusted R^2^ values and P-values (derived from Pearson correlations, equivalent to simple OLS tests) are likewise displayed only when p < 0.1. Error bars denote ± standard error of the mean; colours denote different populations.

**Figure 8:**
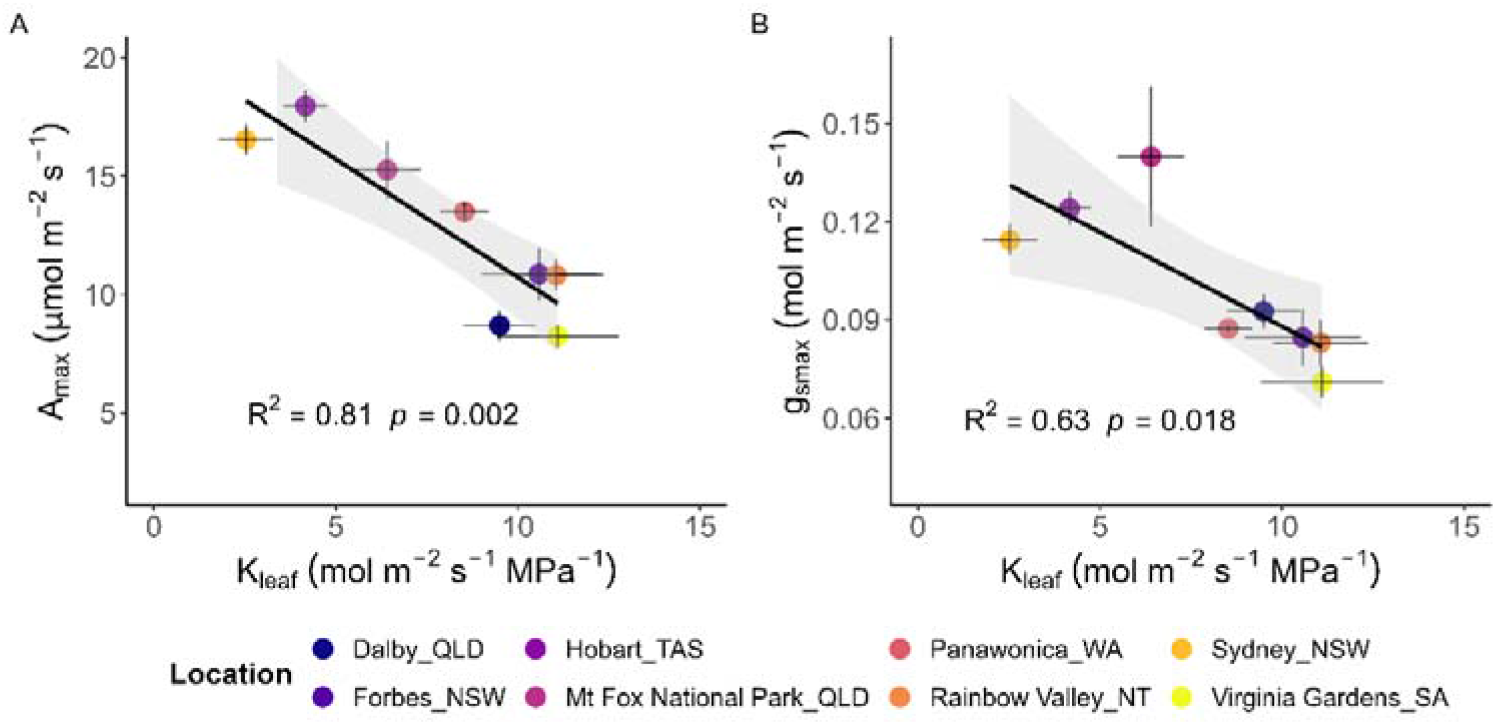
Linear regressions examining the relationships between leaf hydraulic conductance and leaf gas exchange across populations of *T. triandra*. Correlations between leaf hydraulic conductance (K_leaf_) and (A) maximum photosynthetic rate (A_max_) and (B) maximum stomatal conductance (g_smax_). Solid lines and shaded 95% confidence bands are displayed only for statistically significant relationships (P < 0.1). Adjusted R^2^ values and P-values (derived from Pearson correlations, equivalent to simple OLS tests) are likewise displayed only when p < 0.1. Error bars denote ± standard error of the mean; colours denote different populations.

## Discussion

Our study of key hydraulic, anatomical, and photosynthetic traits measured in eight populations of *T. triandra* selected from diverse climatic regions across Australia revealed contrasting trait syndromes associated with climate-of-origin precipitation and temperature. Our results challenge commonly assumed trait trade-offs, suggesting a more nuanced coordination of hydraulic function and gas exchange traits across populations of this widespread grass species. These findings provide insight into how diverse hydraulic strategies emerge within a single species and highlight key traits that may enhance drought resistance and resilience under future climates.

### High hydraulic conductance and embolism resistance in warmer climates

Consistent with our second hypothesis, plants from warm sites exhibited generally higher hydraulic conductance, aligning with previous findings in woody angiosperms (Nardini & Luglio, 2014; Larter *et al*., 2017; Peters *et al*., 2021). Plants growing in hot, high-VPD environments may require greater hydraulic capacity to match elevated evaporative demand under such conditions (Brodribb & Jordan, 2008; Osborne & Sack, 2012; Scoffoni *et al*., 2016). Beyond maintaining plant water balance, higher K_leaf_ also facilitates greater leaf transpirational cooling in hot environments (Blonder *et al*., 2023). Notably, plants from warmer sites also exhibited more negative P_50_, suggesting increased embolism resistance is adaptive where high evaporative demand and increased transpiration generate strongly negative xylem water potentials. This pattern of increasing embolism resistance with increasing temperature has been documented in other herbaceous (Ocheltree *et al*., 2016; Huang *et al*., 2024, p. 202) and woody species (Trueba *et al*., 2017; Guan *et al*., 2023), as well as in different provenances of the tree species *Corymbia calophylla* (Blackman *et al*., 2017). However, temperature-P_50_ relationships in woody species vary considerably across studies, with some finding no relationship (Larter *et al*., 2017; Li *et al*., 2018; López *et al*., 2021; Peters *et al*., 2021) or even negative correlations on a global scale (Nardini & Luglio, 2014). This variation may partly reflect confounding effects of temperature and precipitation, making it difficult to isolate the independent effects of each climate variable. In our study, low collinearity between these factors allowed us to separate their independent effects. Additionally, variation in temperature-P_50_ relationships may reflect different selective pressures operating at temperature extremes, whereby high embolism resistance protects against water stress-induced cavitation in hot environments and against freeze-thaw induced embolism in very cold environments (Davis *et al*., 1999; Pittermann & Sperry, 2006; Hartill *et al*., 2023). Since the *T. triandra* populations in our study originated from regions that rarely see severe freezing events during the growing season, the observed increase in embolism resistance with increasing temperature likely reflects adaptations to hydraulic stress due to high VPD in warm climates rather than to freeze-thaw embolism tolerance in cooler climates.

Plants from warmer sites exhibited larger minor vein vessels (whereas precipitation was unrelated to anatomical traits). These patterns accord with our second hypothesis and mirror observations in woody species where temperature—but not precipitation—predicted vessel diameter (Morris *et al*., 2018; Olson *et al*., 2018), suggesting stronger temperature-driven selection on leaf xylem architecture. Larger vessels may be advantageous in warmer environments with high transpirational demand, as small increases in conduit size can yield large increases in vascular hydraulic conductance according to the Hagen-Poiseuille relationship (Sperry *et al*., 2006). However, we did not observe a corresponding increase in total K_leaf_ with vessel diameter, likely because total K_leaf_ integrates conductance across both xylem and outside-xylem pathways, with the latter typically representing the dominant hydraulic bottleneck (Scoffoni *et al*., 2023).

### Conservative water use and efficient water transport in dry environments

Our finding that leaf P_50_ was not related to precipitation did not accord with our first hypothesis or numerous cross-species studies that have reported increased embolism resistance in drier environments (Nardini & Luglio, 2014; Blackman *et al*., 2014; Bourne *et al*., 2017; Li *et al*., 2018; Volaire *et al*., 2018; Yao *et al*., 2021). However, intraspecific studies frequently reveal weak or absent correlations between P_50_ and precipitation (Martínez-Vilalta *et al*., 2009; Bayramzadeh *et al*., 2011; Lamy *et al*., 2014; Blackman *et al*., 2017; López *et al*., 2021), suggesting that populations from drier climates make adjustments in other traits (e.g., sapwood-to-leaf area ratio (Huber value), stem hydraulic conductivity) that help maintain water potentials within safe limits. Nevertheless, the lack of a P_50_-precipitation relationship in *T. triandra* parallels patterns observed in nine C_4_ grass species (Ocheltree *et al*., 2016) and may reflect differences in drought response strategies between woody plants and grasses. Woody species (especially evergreens) generally maintain the same conductive tissues through successive periods of water deficit, exposing them to repeated embolism risk. In contrast, grasses such as *T. triandra* can senesce above-ground organs during drought and resprout via basal meristems once drought is alleviated (Volaire *et al*., 2009; VanderWeide *et al*., 2014), reducing the need to maintain embolism-resistant tissues.

Although we did not observe a significant correlation between P_50_ and climate of origin precipitation, populations from drier sites closed their stomata at less negative water potentials during drought than populations from wetter sites. This earlier stomatal closure in effect increased the stomatal safety margin for drier-site populations (Fig. 2E), indicating a more conservative water use strategy, favouring drought avoidance over tolerance. The observation of less negative g_s90_ values in dry-site *T. triandra* populations aligns with findings among closely-related *Caragana* shrub species (Yao *et al*., 2021) but contrasts with patterns documented in Australian trees, in which stomatal closure generally occurs at more negative water potentials in drier climates (Bourne *et al*., 2017; Li *et al*., 2018). Our observation of less negative Ψ_TLP_ in dry-site populations is opposite in sign to the relationship reported from several comparative studies across predominantly woody taxa (Mitchell & O’Grady, 2015; Bourne *et al*., 2017) yet this discrepancy aligns with evidence that relationships between turgor loss and climate can differ between species (López *et al*., 2021) and are weak within grasses (Ocheltree *et al*., 2016). .

*T. triandra* plants from drier climates exhibited higher K_leaf_ than those from wetter climates. While this pattern runs counter to general trends in woody species (Nardini & Luglio, 2014) it is consistent with findings in closely related *Caragana* species (Yao *et al*., 2021) and within *Pinus canariensis* (López *et al*., 2016). Higher K_leaf_ in dry-site populations may enhance the capacity of shallow-rooted grasses to capitalise on brief rainfall pulses in otherwise water-limited environments. Plants from dry sites also exhibited significantly lower rates of g_smax_, which may be an adaptation to mitigate water loss from high VPD and low water availability. Taken together, these patterns suggest that *T. triandra* populations from drier climates employ a drought-avoidance strategy characterised by earlier turgor loss and stomatal closure, higher K_leaf_, and reduced operating stomatal conductance—traits that together minimise exposure to critically low water potentials while maximising the capacity to exploit transient increases in soil moisture.

### Coordination among hydraulic traits

Although empirical evidence is mixed, in the sapwood of woody species it is often assumed there is a trade-off between xylem hydraulic safety and efficiency, whereby more conductive wider vessels are inherently more vulnerable to cavitation (Hacke *et al*., 2006; Gleason *et al*., 2016). The same principle for a trade-off exists in leaves and has been observed in some groups of species (Nardini & Luglio, 2014; Blackman *et al*., 2024) but not others (Chen *et al*., 2009; Hao *et al*., 2013). Here, P_50_ and K_leaf_ were unrelated across populations (Fig. 5A); moreover, plants from warmer sites exhibited both high embolism resistance and high K_leaf_.

Variation in K_leaf_ was also unrelated to the diameter of major and minor vein vessels and was negatively correlated with minor vein density. This was unexpected considering that high K_leaf_ is typically associated with wide vessels (Sperry *et al*., 2006; Zanne *et al*., 2010) and high vein density (Brodribb *et al*., 2007). Our observations suggest that leaf hydraulic function in *T. triandra* may be primarily determined by outside-xylem factors rather than xylem vessel anatomy. In leaves, outside-xylem pathways have been shown to contribute substantially to hydraulic conductance and influence leaf hydraulic decline during drought (Scoffoni *et al*., 2023). Because reductions in outside-xylem conductance (K_ox_) are generally recoverable (Scoffoni et al., 2014; Zhang et al., 2016), while xylem embolism is not (Skelton *et al*., 2017; Johnson *et al*., 2018), loss of K_ox_ may act as a protective buffer, decoupling leaf hydraulic safety from efficiency. Consistent with this mechanism, K_leaf_ has been reported to decline well before the onset of xylem embolism in other herbaceous species (Corso *et al*., 2020; Yao *et al*., 2021), including *T. triandra* (Jacob *et al*., 2022).

Turgor loss point (Ψ_TLP_) was strongly coordinated with g_s90_ (Fig. 7E), reinforcing loss of cell turgor as a key driver of stomatal closure (Bartlett *et al*., 2012, 2014). The combination of low g_smax_ and early loss of turgor in *T. triandra* plants from dry sites suggests a coordinated strategy to conserve water under water limiting conditions. P_50_ and g_s90_ were unrelated, contrasting with the idea that stomatal downregulation under drought is caused by embolism-induced reductions in K_leaf_ (Nardini & Salleo, 2000; Brodribb & Holbrook, 2003; Bartlett *et al*., 2016). Instead, our results are consistent with recent studies on herbaceous species which also observed no coordination between xylem embolism and stomatal closure (Corso *et al*., 2020; Jacob *et al*., 2022).

### Hydraulic constraints on leaf gas exchange relaxed by C_4_ metabolism

K_leaf_ was negatively correlated with maximum stomatal conductance (g_smax_) and photosynthetic capacity (A_max_). While this contrasts with the positive coordination between hydraulic and photosynthetic traits sometimes reported in C_3_ species (Franks, 2006; Brodribb & Holbrook, 2006; Mitchell *et al*., 2014; Xiong *et al*., 2018), it is consistent with work on other C_4_ grasses, in which K_leaf_ and leaf gas exchange traits were found to be unrelated (Ocheltree *et al*., 2016; Zhou *et al*., 2020; Pathare *et al*., 2020; Baird *et al*., 2025). It seems that C_4_ grasses may not require high K_leaf_ to sustain photosynthetic rates (unlike in C_3_ species), as their carbon-concentrating mechanism allows high photosynthesis despite lower g_s_ and less negative operating Ψ_leaf_ (Taylor *et al*., 2011; Zhou *et al*., 2020; Baird *et al*., 2025). Additionally, traits unique to C_4_ metabolism — such as variation in bundle sheath leakiness — may alter the balance between carbon gain and water movement. Species with low bundle sheath leakiness have the ability to maintain high photosynthetic rates even when the hydraulic conductance is low (Fravolini, 2002). The decoupling may also reflect the pulse-driven environments many C_4_ grasses occupy, where the need for maintaining hydraulic safety during dry periods and rapidly maximising carbon gain after rainfall further weaken correlations between K_leaf_ and A_max_.

### Conclusion

Hydraulic strategies of the widespread perennial grass *T. triandra* highlight the complex interplay between hydraulic traits, stomatal regulation and xylem anatomy in shaping climate adaptation. Our findings indicate that populations from drier climates exhibit traits associated with conservative water-use and drought avoidance, displaying lower stomatal conductance, less negative g_s90_, less negative Ψ_TLP_ and larger stomatal safety margins than wetter-site populations. By contrast, plants from warmer climates exhibited a combination of high hydraulic conductance and high embolism resistance, enabling rapid water transport to match periods of high evaporative demand, while reducing the risk of hydraulic failure. Our results reveal substantial within-species variation in hydraulic traits and provide a more nuanced understanding of the mechanisms enabling success in dry and hot environments that may help inform vegetation models aimed at predicting plant responses to intensifying drought and warming in grassland ecosystems.

## Acknowledgements

We thank Dr Alan Humphries from the Australian Pastures Genebank for advice on germinating *T. triandra* and for providing seed sources that helped make this project possible. VKJ, IJW, CJB, EF, ACL TB and ES acknowledge support from the Australian Research Council (CE200100015).

## Author contributions

VJ and IJW led project design, experimental work, and manuscript preparation, with strong support from CJB and BC. ACL, TB, and ES assisted with data collection. All authors discussed results, contributed ideas and text to multiple versions of the manuscript, and approved the final version.

