## Supplementary for "Linking drought strategy to climate of origin in a widespread C4 grass: hydraulic traits, xylem anatomy and stomatal behaviour in *Themeda triandra*"

### Supplementary figures and tables

Table S1: Bivariate analysis of traits vs climate

| **Trait** | **Predictor** | **P value** | **Marginal R^2^** | **Conditional R^2^** |
| --- | --- | --- | --- | --- |
| P_50_ | Temperature | **0.038** | **0.295** | **0.434** |
| P_50_ | Precipitation | 0.696 | 0.013 | 0.485 |
| ψ_TLP_ | Temperature | 0.703 | 0.012 | 0.442 |
| ψ_TLP_ | Precipitation | **0.027** | **0.268** | **0.395** |
| g_s90_ | Temperature | 0.303 | 0.037 | NA |
| g_s90_ | Precipitation | **0.013** | **0.619** | NA |
| SSM | Temperature | 0.268 | 0.109 | 0.526 |
| SSM | Precipitation | 0.126 | 0.181 | 0.517 |
| K_leaf_ | Temperature | 0.239 | 0.106 | 0.360 |
| K_leaf_ | Precipitation | 0.128 | 0.159 | 0.346 |
| A_max_ | Temperature | 0.436 | 0.062 | 0.698 |
| A_max_ | Precipitation | 0.229 | 0.141 | 0.691 |
| g_smax_ | Temperature | 0.461 | 0.037 | 0.376 |
| g_smax_ | Precipitation | **0.047** | **0.211** | **0.344** |
| Major Vessel xylem diameter | Temperature | 0.327 | 0.116 | 0.816 |
| Major Vessel xylem diameter | Precipitation | 0.348 | 0.101 | 0.815 |
| Minor Vessel xylem diameter | Temperature | **0.009** | **0.458** | **0.590** |
| Minor Vessel xylem diameter | Precipitation | 0.918 | 0.001 | 0.629 |
| Major Vein Density | Temperature | 0.251 | 0.070 | 0.221 |
| Major Vein Density | Precipitation | 0.473 | 0.028 | 0.232 |
| Minor Vein Density | Temperature | **0.052** | **0.304** | **0.593** |
| Minor Vein Density | Precipitation | 0.604 | 0.026 | 0.615 |
| **Moisture Index** | | | | |
| **Trait** | **Predictor** | **P value** | **Marginal R^2^** | **Conditional R^2^** |
| P_50_ | Moisture Index | 0.339 | 0.079 | 0.476 |
| ψ_TLP_ | Moisture Index | 0.863 | 0.002 | 0.442 |
| g_s90_ | Moisture Index | 0.967 | -0.166 | NA |
| SSM | Moisture Index | 0.372 | 0.074 | 0.532 |
| K_leaf_ | Moisture Index | **0.067** | **0.203** | **0.348** |
| A_max_ | Moisture Index | 0.324 | 0.097 | 0.695 |
| g_smax_ | Moisture Index | 0.388 | 0.050 | 0.374 |
| Major Vessel xylem diameter | Moisture Index | 0.596 | 0.035 | 0.819 |
| Minor Vessel xylem diameter | Moisture Index | 0.102 | 0.232 | 0.610 |
| Major Vein Density | Moisture Index | 0.424 | 0.036 | 0.226 |
| Minor Vein Density | Moisture Index | **0.049** | **0.307** | **0.593** |

Table S2: Temperature and Precipitation variables of populations used in experiment. Column names : MI = Moisture Index, MAT = Mean Annual Temperature, MAP = Mean Annual Precipitation, MTWQ = Summer Temperature, PWQ = Summer Precipitation. Populations are ordered by increasing MAP.

| **Location** | **latitude** | **longitude** | **MI** | **MAT** | **MAP** | **MTWQ** | **PWQ** |
| --- | --- | --- | --- | --- | --- | --- | --- |
| Rainbow Valley | -24.2376 | 133.5536 | 0.1613 | 22.05 | 281.1 | 29.55 | 116.4 |
| Pannawonica | -21.6449 | 116.3233 | 0.1937 | 26.85 | 387.0 | 32.05 | 213.8 |
| Forbes | -33.4076 | 147.9655 | 0.3745 | 16.95 | 489.9 | 24.55 | 127.1 |
| Hobart | -42.8651 | 147.331 | 0.6337 | 12.45 | 536.0 | 16.25 | 133.6 |
| Dalby | -27.1149 | 151.1818 | 0.4297 | 19.55 | 627.2 | 25.15 | 250.4 |
| Virginia | -34.8539 | 138.6206 | 0.6480 | 16.15 | 667.6 | 21.05 | 75.1 |
| Mt Fox National Park | -19.0013 | 145.4731 | 0.4501 | 22.65 | 711.5 | 26.35 | 435.9 |
| Sydney | -33.6881 | 151.0624 | 1.0404 | 16.55 | 1129.1 | 21.45 | 333.9 |

Table S3: *Principal component analysis (varimax-rotated) for ten CHELSA climate variables describing temperature and precipitation at the eight populations. Loadings are shown for the first two rotated PCs; bold values indicate the strongest loadings on each axis, and the variables subsequently selected for analyses.*

| Variable | PC1 | PC2 |
| --- | --- | --- |
| Moisture Index | 0.224578 | 0.371783 |
| *Mean Annual Temperature* | -0.40465 | 0.119154 |
| *Summer Temperature* | **-0.47485** | 0.009087 |
| *Mean Annual Precipitation* | 0.039343 | 0.550788 |
| Precipitation Seasonality | -0.12575 | 0.174117 |
| Precipitation of the Driest Quarter | 0.060642 | 0.242834 |
| Summer Precipitation | -0.26936 | **0.626561** |
| Temperature of the warmest month | -0.45144 | -0.04577 |
| Annual temperature range | -0.50852 | -0.21675 |
| Temperature of the Driest Quarter | 0.066008 | -0.11484 |

Table S4: Methodology used for quantifying xylem anatomical traits across populations of T. triandra. *Measurements were performed on transverse leaf sections stained with* Safranin O and Fast Green*. Vessel diameter was measured as the maximum distance across the lumen of metaxylem vessels for major vessels (blue arrow) and minor vessels (red arrow). Major vessel metaxylem wall thickness was quantified as the distance from the inner to outer wall boundary of metaxylem vessels (orange line). For each sample, two vessels were measured from two randomly selected vascular bundles to obtain mean values per individual.* M*ajor and minor veins were individually counted on two leaf sections per plant and vein density was quantified as the total number of veins per unit leaf size (cm^-1^). Measurements were taken from digital microscope images, at a 100x magnification, using image J to normalise pixel information into um scale to quantify distances. The observations were recorded using the ‘polygon selection,’ ‘Straight’ and ‘Multi-point’ tools to log areas, diameters/lengths and count elements, respectively.*

| Metric | Example Image |
| --- | --- |
| Major vessel diameter  Major vessel metaxylem wall thickness  Minor vessel diameter | 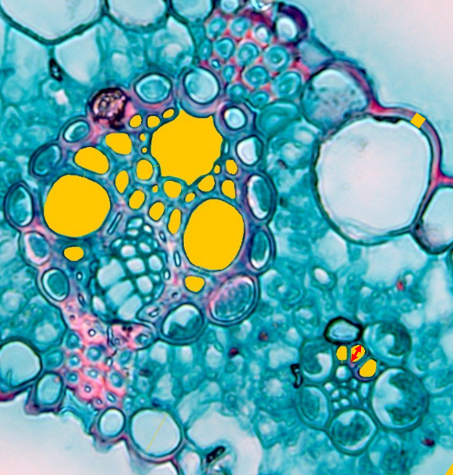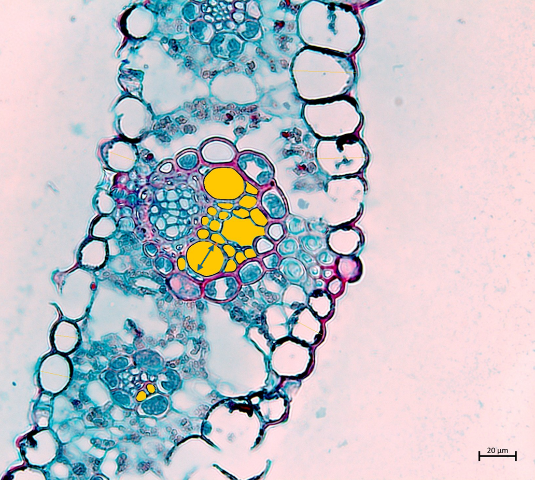 |
| Major and minor veins counted (yellow dots).  White line surrounding leaf border used to calculate length of leaf section.  Vein density quantified as vein number (No.) per leaf width (cm^-1^) | 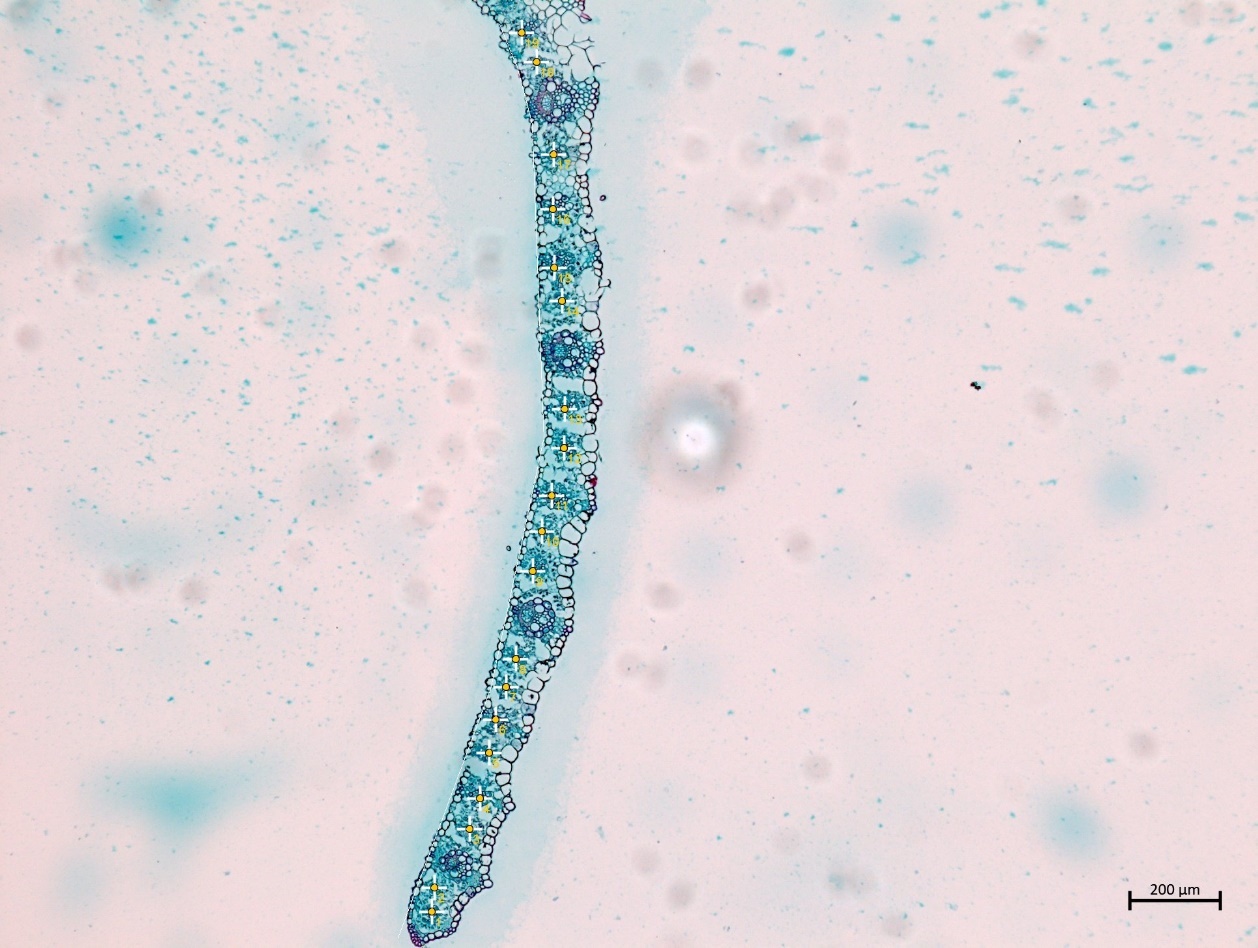 |

Table S5: *Population -level mean values of hydraulic, anatomical, and gas-exchange traits in Themeda triandra.*

| Accession | P_50_ | Ψ_TLP_, | g_s90_ | SSM | K_leaf_ | Major vein xylem diameter | Major vein Density | Minor vein xylem diameter | Minor vein density | Vessel wall thickness | A_max_ | g_smax_ |
| --- | --- | --- | --- | --- | --- | --- | --- | --- | --- | --- | --- | --- |
| Dalby | 3.378 | 1.588 | 4.383 | -1.006 | 9.495 | 17.560 | 25.945 | 4.412 | 103.983 | 1.701 | 8.698 | 0.093 |
| Forbes | 4.677 | 1.534 | 3.926 | 0.751 | 10.575 | 21.099 | 28.678 | 4.770 | 93.058 | 1.670 | 10.870 | 0.085 |
| Hobart | 2.414 | 1.663 | 3.931 | -1.517 | 4.161 | 13.621 | 24.464 | 3.388 | 144.164 | 1.821 | 17.974 | 0.124 |
| Mount Fox | 3.897 | 1.857 | 5.437 | -1.540 | 6.403 | 17.269 | 24.708 | 3.966 | 107.115 | 1.965 | 15.272 | 0.140 |
| Pannawonica | 5.128 | 1.585 | 4.507 | 0.621 | 8.528 | 24.271 | 17.404 | 6.201 | 99.626 | 1.820 | 13.511 | 0.087 |
| Rainbow Valley | 4.808 | 1.735 | 4.581 | 0.227 | 11.057 | 17.922 | 27.084 | 4.598 | 102.030 | 2.121 | 10.829 | 0.083 |
| Sydney | 4.182 | 1.841 | 4.815 | -0.633 | 2.527 | 17.664 | 26.386 | 4.043 | 137.072 | 1.740 | 16.552 | 0.114 |
| Virginia | 4.596 | 1.168 | 3.986 | 0.610 | 11.095 | 25.619 | 27.252 | 4.010 | 101.424 | 2.381 | 8.242 | 0.071 |

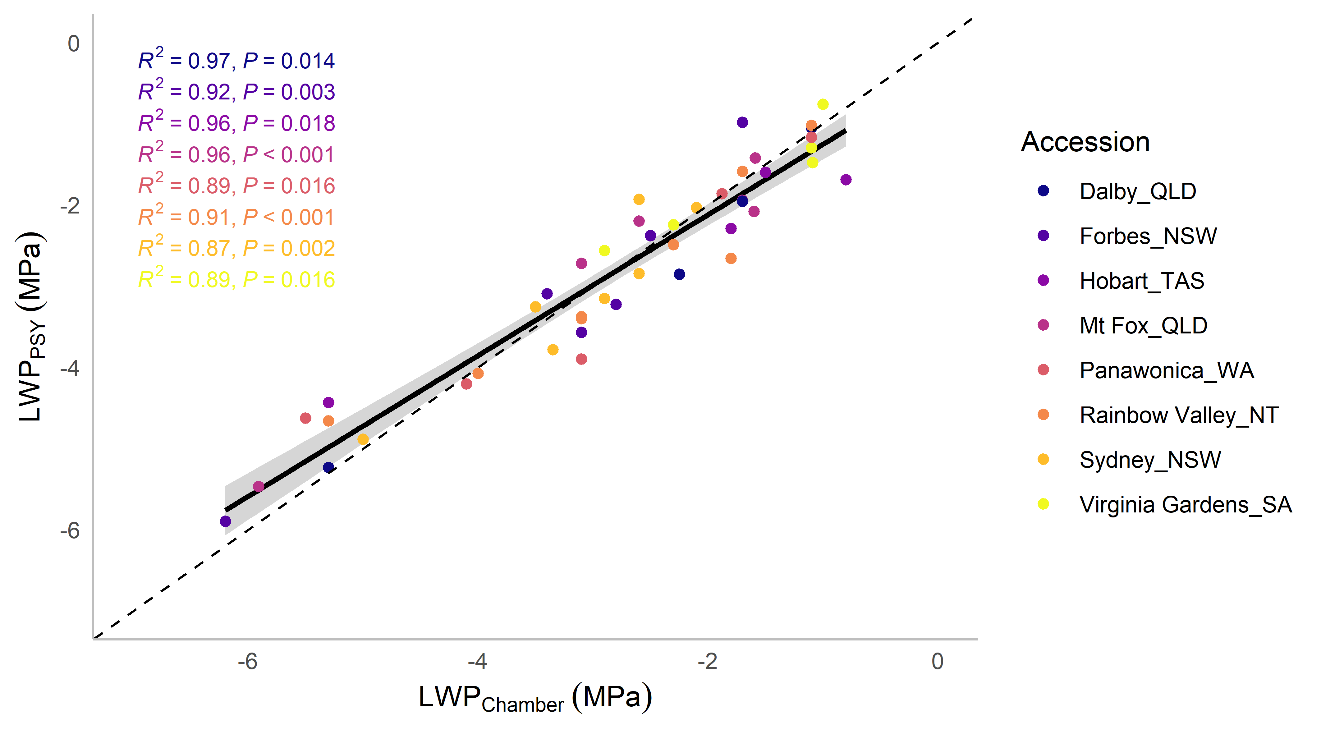

**FIGURE S1**. Psychrometer leaf water potential (LWP) validation, model output of the relationship between LWP measured using a pressure chamber (LWP_Chamber_) and a leaf psychrometer (LWP_PSY_) for the different populations.

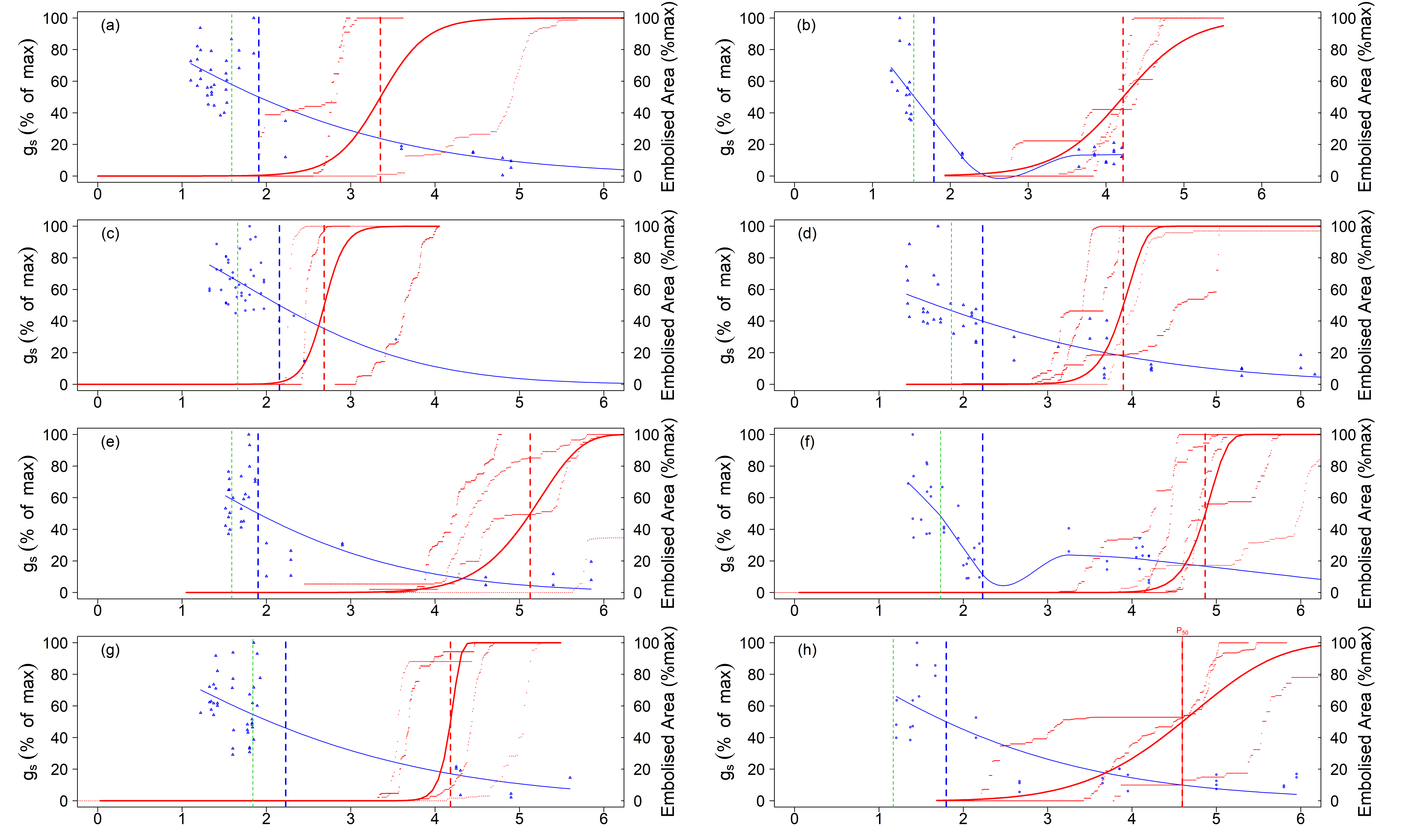

**Figure S2: Percentage loss of conductivity curves for each population:** Relationships between leaf water potential (-MPa), stomatal conductance (g_s_; mol H_2_O m^-2^ s^-1^), and % xylem embolism for (a) Dalby, (b) Forbes, (c) Hobart, (d) Mount-Fox, (e) Pannawonica, (F) Rainbow Valley, (g) Sydney and (h) Virginia (n=3-4). Blue triangles represent g_s_ values with the solid blue line representing model output decline in g_s_ with the vertical dashed blue line indicating the point at which stomata are 50% closed or P_gs50_. The solid red line represents percentage of embolism, modelled using all replicate curves for each species, and the vertical dashed red line indicates the point at which 50% of leaf xylem was embolised (P_X50_). Leaf water potential at turgor loss (ψ_TLP_) is indicated by green vertical dashed lines.
